# Human *HPSE2* gene transfer ameliorates bladder pathophysiology in a mutant mouse model of urofacial syndrome

**DOI:** 10.1101/2023.07.03.547034

**Authors:** Filipa M. Lopes, Celine Grenier, Benjamin W. Jarvis, Sara Al Mahdy, Adrian Lène-McKay, Alison M. Gurney, William G. Newman, Simon N. Waddington, Adrian S. Woolf, Neil A. Roberts

## Abstract

Rare early onset lower urinary tract disorders include defects of functional maturation of the bladder. Current treatments do not target the primary pathobiology of these diseases. Some have a monogenic basis, such as urofacial, or Ochoa, syndrome (UFS). Here, the bladder does not empty fully because of incomplete relaxation of its outflow tract, and subsequent urosepsis can cause kidney failure. UFS is associated with biallelic variants of *HPSE2*, encoding heparanase-2. This protein is detected in pelvic ganglia, autonomic relay stations that innervate the bladder and control voiding. Bladder outflow tracts of *Hpse2* mutant mice display impaired neurogenic relaxation. We hypothesized that *HPSE2* gene transfer soon after birth would ameliorate this defect and explored an adeno-associated viral (*AAV*) vector-based approach. *AAV9/HPSE2,* carrying human *HPSE2* driven by *CAG*, was administered intravenously into neonatal mice. In the third postnatal week, transgene transduction and expression were sought, and *ex vivo* myography was undertaken to measure bladder function. In mice administered *AAV9/HPSE2*, the viral genome was detected in pelvic ganglia. Human *HPSE2* was expressed and heparanase-2 became detectable in pelvic ganglia of treated mutant mice. On autopsy, wild-type mice had empty bladders whereas bladders were uniformly distended in mutant mice, a defect ameliorated by *AAV9/HPSE2* treatment. Therapeutically, *AAV9/HPSE2* significantly ameliorated impaired neurogenic relaxation of *Hpse2* mutant bladder outflow tracts. Impaired neurogenic contractility of mutant detrusor smooth muscle was also significantly improved. These results constitute first steps towards curing UFS, a clinically devastating genetic disease featuring a bladder autonomic neuropathy.

**Summary:** In the first gene therapy for genetic bladder disease, we cured autonomic neurons using AAV-mediated gene delivery in a mouse model of urofacial syndrome.

## INTRODUCTION

Rare early onset lower urinary tract (REOLUT) disorders comprise not only gross anatomical malformations but also primary defects of functional maturation (Woolf *et al*, 2019). Although individually infrequent, these diseases are together a common cause of kidney failure in children and young adults (Harambat *et al*, 2012; Woolf, 2022; Pepper & Trompeter, 2022). REOLUT disorders can also have negative impacts on the self-esteem, education, and socialization of affected individuals (Pepper & Trompeter 2022; Hankinson *et al*, 2014). They have diverse phenotypes including: ureter malformations, such as megaureter; bladder malformations, such as exstrophy; and bladder outflow obstruction caused either by anatomical obstruction, as in urethral valves, or functional impairment of voiding without anatomical obstruction. The latter scenario occurs in urofacial, or Ochoa, syndrome (UFS) (Ochoa, 2004).

During healthy urinary voiding, the bladder outflow tract, comprising smooth muscle around the section of the urethra nearest the bladder, dilates while detrusor smooth muscle in the bladder body contracts. Conversely, in UFS, voiding is incomplete because of dyssynergia in which the outflow tract fails to fully dilate (Ochoa, 2004), and subsequent accumulation of urine in the LUT predisposes to urosepsis (Ochoa, 2004; Osorio *et al*, 2021). UFS is an autosomal recessive disease and around half of families studied genetically carry biallelic variants in *HPSE2* (Daly *et al*, 2010; Pang *et al*, 2010; Newman *et al*, 2023). Most are frameshift or stop variants, most likely null alleles, although missense changes, triplication and deletions have also been reported (Beaman *et al*, 2022). *HPSE2* codes for heparanase-2 (McKenzie *et al*, 2000), also known as Hpa2, which inhibits endoglycosidase activity of the classic heparanase (Levy-Adam *et al*, 2010). Although subject to secretion (Levy-Adam *et al*, 2010; Beaman *et al*, 2022), heparanase-2 has also been detected in the perinuclear membrane (Margulis *et al*, 2021), suggesting yet-to-be defined functions there. The biology of heparanase-2 has been most studied in relation to oncology. For example, in head and neck cancers, heparanase-2 has anti-tumour effects, enhancing epithelial characteristics and attenuating metastasis (Gross-Cohen *et al*, 2021).

The autonomic nervous system controls voiding of the healthy bladder (Keast *et al*, 2015). In fetal humans and mice, heparanase-2 is immunodetected in bladder nerves (Stuart *et al*, 2013; Stuart *et al*, 2015). The protein is also present in pelvic ganglia near the bladder (Stuart *et al,* 2015; Roberts *et al*, 2019). These ganglia contain neural cell bodies that send autonomic effector axons into the bladder body and its outflow tract, with this neural network maturing after birth in rodents (Keast *et al*, 2015; Roberts *et al*, 2019). Mice carrying biallelic gene-trap mutations of *Hpse2* have bladders that fail to fully void despite the absence of anatomical obstruction within the urethral lumen (Stuart *et al*, 2015; Guo *et al*, 2015). Although *Hpse2* mutant mice do have pelvic ganglia, autonomic nerves implicated in voiding are abnormally patterned (Roberts *et al*, 2019). *Ex vivo* physiology experiments with *Hpse2* mutant juvenile mice demonstrate impaired neurogenic bladder outflow tract relaxation (Manak *et al*, 2020), an observation broadly consistent with the functional bladder outflow obstruction reported in people with UFS (Ochoa, 2004). Therefore, while not excluding additional aberrations, such as a central nervous system defect (Ochoa, 2004), much evidence points to UFS featuring a genetic autonomic neuropathy affecting the bladder (Roberts & Woolf, 2020).

Recently, gene therapy has been used to treat animal models and human patients with previously incurable genetic diseases. Adeno associated virus 9 (*AAV9*) has been used as a vector in successful gene therapy for spinal muscular atrophy, an early onset genetic neural disease (Mendell *et al*, 2017). Here, we hypothesized that human *HPSE2* gene therapy would restore autonomic nerve function in the bladder. Given that UFS is a neural disease, we elected to use *AAV*2/9 (simplified to *AAV9*) vector, which transduces diverse neural tissues in mice (Foust *et al*, 2009), although uptake into the pelvic ganglia has not been reported. We administered *AAV9* vector carrying human *HPSE2* to neonatal mice via the temporal vein, in the first day after birth when bladder neural circuitry is maturing. Several weeks later, evidence of transgene transduction and expression were sought and therapeutic effects on neuropathic dysfunction were investigated in smooth muscle relaxation of the bladder outflow tract and smooth muscle contractility of the bladder body.

## RESULTS

### Administration of AAV9/HPSE2 to WT mice

We undertook exploratory experiments in *WT* mice, primarily to determine whether the viral vector was capable of transducing pelvic ganglia after neonatal intravenous injection. We also wished to determine whether administration of *AAV9/HPSE2* was compatible with normal postnatal growth and general health. Given that human gene therapy with *AAV9* vectors have been associated with a specific side-effect, liver damage (Mendell *et al*, 2017), and the recent implication of AAV2 in community acquired hepatitis (Ho *et al,* 2023), we also examined mouse livers at the end of the experiment. To determine efficacy of transduction, neonatal *WT* mice were administered the *AAV9*/*HPSE2* vector, and they were followed until they were young adults (Figure 1a). Previous studies administering *AAV9* carrying reporter genes to neonatal mice have used single doses of up to 10^12^ genome copies (Foust *et al*, 2009; Buckinx *et al*, 2016). In the current experiments, because the *AAV9*/*HPSE2* vector was yet to be tested *in vivo*, we took a cautious approach, assessing single doses of 2x10^10^ or 1x10^11^ genome copies. *WT* neonatal mice administered either dose gained body weight in a similar manner to *WT* mice injected with vehicle only (Figure 1b). Moreover, throughout the five-week observation period, *AAV9/HPSE2* administered mice displayed normal general appearances and behaviour (e.g., condition of skin and fur, ambulation, grooming, and feeding). As outlined in the *Introduction*, evidence points to UFS being an autonomic neuropathy of the bladder. Therefore, we determined whether the *AAV9/HPSE2* vector had targeted pelvic ganglia that flank the base of the mouse bladder (Keast *et al*, 2015; Roberts *et al*, 2019).

**Figure 1.**
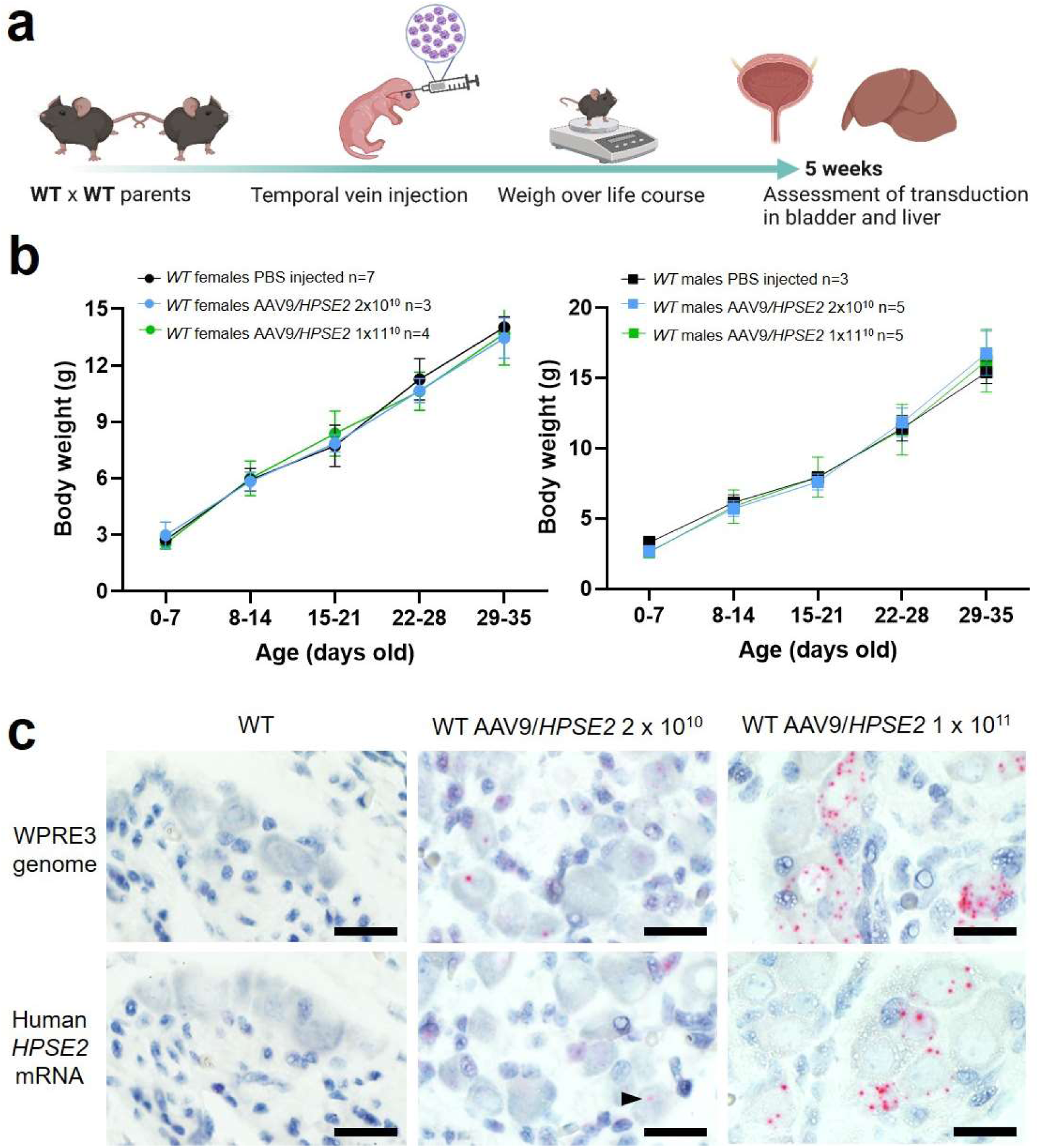
Administration of *AAV9*/*HPSE2* to neonatal *WT* mice. **a.** Graphic of study design. The *AAV9/HPSE2* vector genome consisting of flanking AAV2 ITR sequences, the ubiquitous CAG promoter, human *HPSE2* coding sequence, and the WPRE3 sequence. A single dose (2x10^10^ or 1x10^11^ genome copies) of *AAV9/HPSE2* was administered to neonates *via* the temporal vein. Another group of mice received vehicle-only injections. Body weights were monitored, and bladders and livers were harvested for histology analyses at five weeks. Graphic created BioRender.com. **b.** Whole body weights (g). No significant differences were found in growth trajectories comparing: 2x10^10^ *AAV9*/*HPSE2* injected mice with vehicle-only controls (two-way ANOVA); 1x10^11^ *AAV9*/*HPSE2* injected compared with vehicle-only injected controls; and lower dose compared with higher dose *AAV9/HPSE2* injected mice. **c.** Basescope^TM^ ISH of pelvic ganglia. Histology sections are counterstained so that nuclei appear blue. Images are representative of ganglia from three mice in each experimental group. *WPRE3* genomic sequence and human *HPSE2* transcripts were not detected in pelvic ganglia of *WT* mice injected with vehicle-only (left panel). In contrast, positive signals (red dots) for each probe were detected in pelvic ganglia of *WT* mice administered either 2x10^10^ (arrow head) or 1x10^11^ *AAV9*/*HPSE2*. For quantification of positive signals, see main *Results* text. Bars are 20 μm.

To seek the transduced vector genome, tissue sections were stained using Basescope^TM^ *in situ hybridisation* (ISH) for the *WPRE3* genomic sequence (Figure 1c, upper panels). The regulatory element was detected in neural cell bodies of ganglia of five-week-old mice that had been administered either dose of *AAV9/HPSE2,* while no signal was detected in ganglia of vehicle-only injected mice. Of 42 ganglion cells imaged from three mice injected with the 2x10^10^ dose, and 71 ganglion cells imaged from three mice injected with the 10^11^ dose, 45% were positive in each group. Here, the definition of positive was at least one red dot in a cell. The transduced cargo was expressed, as indicated by reaction with the human *HPSE2* mRNA probe (Figure 1c). No signal was detected in mice that received vehicle-alone. In contrast, *HPSE2* transcripts were detected in pelvic ganglia of mice that had been administered *AAV9/HPSE2*. Of 44 ganglion cells imaged from three mice injected with the 2x10^10^ dose, 11% were positive; and of 57 ganglion cells imaged from three mice injected with the 10^11^ dose, 40% were positive. Moreover, although not quantified, the intensity of expression within positive cells appeared greater in the higher dose group (Figure 1c).

Liver sections (Figure 2) were assessed for *WPRE3* Basescope^TM^ signals by counting the number of red dots per unit area. The average density measured in sections from three mice injected with 2x10^10^ vector was 587/mm^2^, compared with 835/mm^2^ for 3 mice injected with 2x10^11^ vector, while sections from vehicle-injected mice showed no signal. Assessing *HPSE2* Basescope^TM^ signals, the density averaged 45/mm^2^ for three mice injected with 2x10^10^ vector, 680/mm^2^ for three mice injected with 2x10^11^ vector and zero for vehicle-injected mice. PSR-stained liver sections, imaged with bright-field and polarised light are illustrated in Figure 2b. The birefringence under polarised light, indicating collagen fibres, was measured as the percentage of the field of view it occupied (Figure 2c). There was no significant difference among the three animal groups in the amount of collagen present, suggesting that neonatal administration of *AAV9/HPSE2* was not associated with liver fibrosis when assessed in mouse early adulthood.

**Figure 2.**
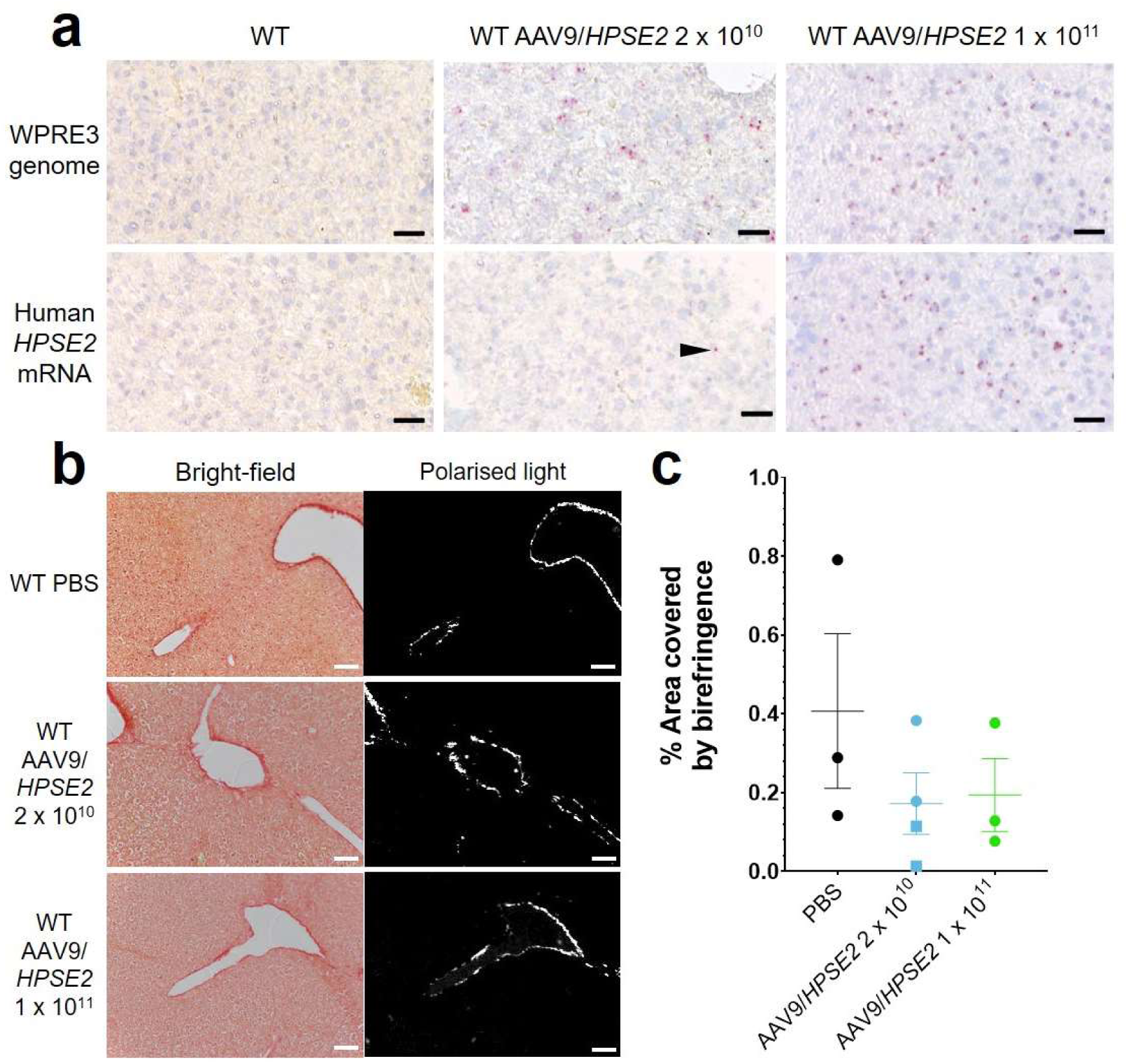
Histology of livers of five week old *WT* mice that had been administered *AAV9*/*HPSE2* as neonates. In each experimental group, livers from three mice were examined, and representative images are shown. **a.** Basescope^TM^ ISH analyses of livers, with positive signals appearing as red dots. Nuclei were counterstained with haematoxylin. Note the absence of signal for *WPRE3*, part of the *AAV9/HPSE2* genomic cargo, in mice administered PBS only. In contrast, *WPRE3* signal was evident in livers of mice that had received a single dose of 2x10^10^ or 1x10^11^ genome copies. Regarding human *HPSE2* transcripts, none were detected in the PBS-only livers. Only sparse signals were noted in livers of mice administered the lower AAV dose (arrow head) but the signal for *HPSE2* was prominent in livers of mice administered the higher dose. See main text for quantification of signals. **b.** PSR staining to seek collagen imaged under direct light (left column) and polarised light (right column). There was no significant difference in extent of birefringence between the three groups. Black bars are 30 μm, white are 200 μm.

### Administration of *AAV9/HPSE2* to *Hpse2 Mut* mice

Having demonstrated the feasibility and general safety of the procedure in *WT* mice, a second set of experiments were undertaken to determine whether administration of *AAV9/HPSE2* to *Mut* mice would ameliorate aspects of their bladder pathophysiology (Figure 3a). Reasoning that the higher dose *AAV9/HPSE2* (1x10^11^ genome copies) had been well-tolerated in our scoping experiments in *WT* mice, we used it here. As expected (Stuart *et al*, 2015), untreated *Mut* mice gained significantly less body weight than sex-matched *WT* mice over the observation period (Figure 3b). Administration of *AAV9/HPSE2* to *Mut* mice did not significantly modify their impaired growth trajectory (Figure 3b). On autopsy, bladders of untreated *Mut* mice appeared distended with urine (Figure 3c and d), consistent with bladder outflow obstruction, and confirming a previous report (Stuart *et al*, 2015). In contrast, only one of nine bladders of *Mut* mice that had been administered *AAV9/HPSE2* appeared distended (Figure 3c and d). The remainder had autopsy bladder appearances similar to those reported for *WT* mice (Stuart *et al*, 2015). Bladder bodies were then isolated, drained of urine, and weighed. The bladder body/whole mouse weight (Figure 3e) of *Mut* mice that had not received *AAV9/HPSE2* was significantly higher than untreated *WT* mice. While values in *AAV9/HPSE2-* administered *Mut* mice (median 0.0016, range 0.0010 to 0.0139) tended to be higher than those of untreated *WT* mice, they were not significantly different from these controls.

**Figure 3.**
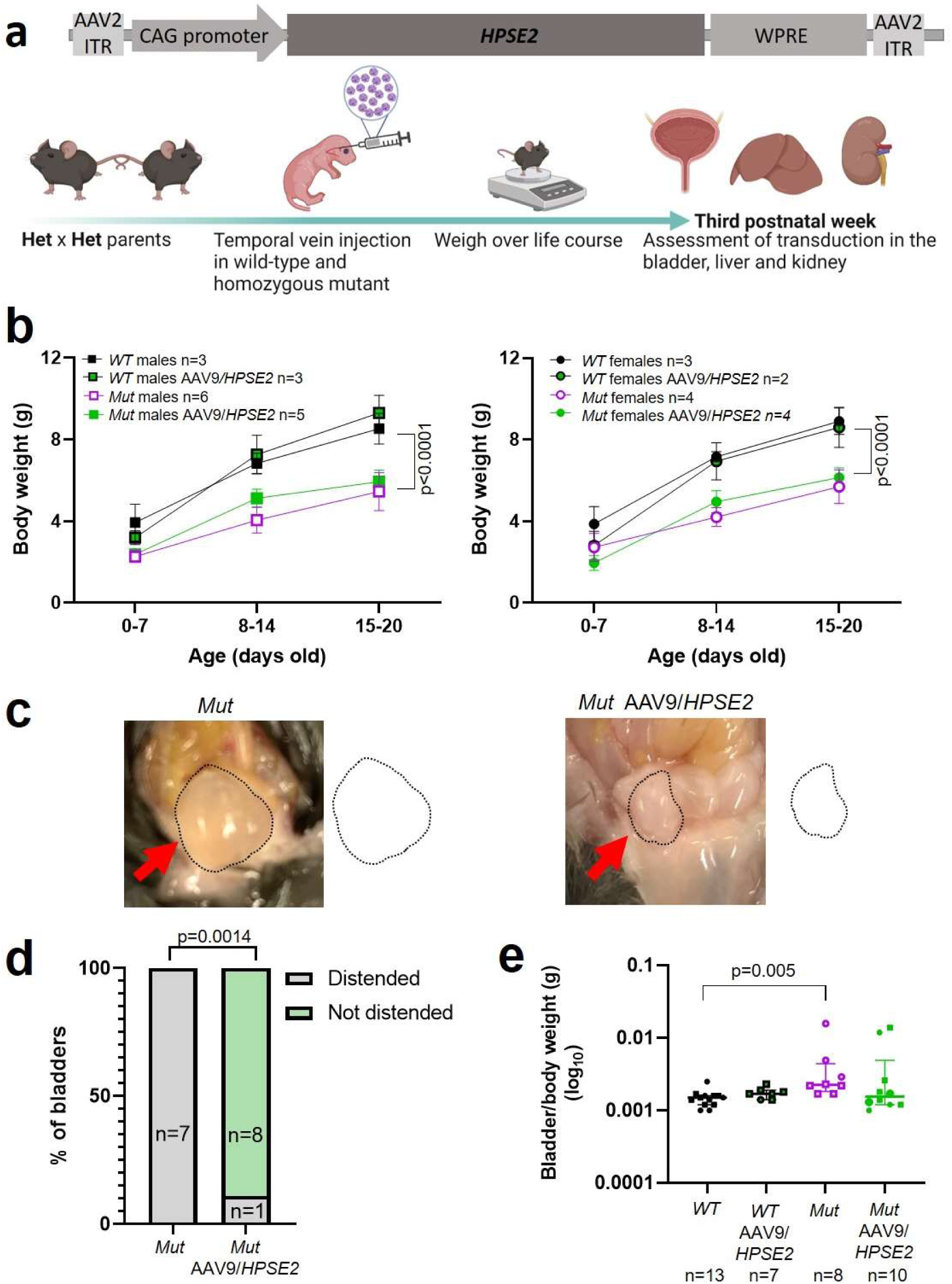
Administration of *AAV9/HPSE2* to neonatal mice. **a.** Graphic of therapy study design, with schematic of *AAV9*/*HPSE2* vector genome. Heterozygous *Hpse2* parents were mated to generate litters and neonates were genotyped, with *WT* and *Mut* offspring used in the study. Some baby mice were not administered the viral vector while others were intravenously administered 1 x 10^11^ *AAV9/HPSE2*. Mice were weighed regularly, and in the third week of postnatal life they were culled and autopsies undertaken to determine whether or not bladders appeared distended with urine. Livers and kidneys were harvested for histology analyses. Bladders were harvested and used either for histology analyses or for *ex vivo* myography. Graphic created BioRender.com. **b**. Body weights (g; mean±SD). Results were analysed with 2-way ANOVA. As expected, body growth was impaired in *Mut* mice that had not received the viral vector compared with sex-matched *WT* mice that did not receive the vector. There was no significant difference in the body growth of *Mut* mice that either had or had not received *AAV9/HPSE2*. **c.** Examples at autopsy of a distended bladder in a *Mut* mouse that had not received the viral vector, and a not-distended bladder in a *Mut* that had been administered *AAV9/HPSE2* as a neonate. Bladder size indicated by dotted line. **d.** Untreated *Mut* mice had distended bladders on autopsy more often than *Mut* mice that had received *AAV9/HPSE2* (Fisher’s exact test). **e.** Untreated *Mut* mice had significantly higher empty bladder/whole body weight ratios than untreated *WT* mice (Kruskal-Wallis test). While viral vector administered *Mut* mice tended to have higher empty bladder/whole body weight ratios than *WT* mice, this was not statistically significant.

Histology sections of pelvic ganglia were reacted with Basescope^TM^ probes (Figure 4). Using the *WPRE3* genomic probe, no signals were detected in either *WT* or *Mut* mice that had not received the viral vector (Figure 4a and c). In contrast, signals for *WPRE3* were detected in *WT* and *Mut* mice administered *AAV9/HPSE2* as neonates (Figure 4b and d). Using the human *HPSE2* probe, no transcripts were detected in ganglia of *WT* and *Mut* mice (Figure 4e and g). In contrast, signals for *HPSE2* were detected in *WT* and *Mut* mice administered *AAV9/HPSE2* as neonates (Figure 4f and h). In sections from *AAV9/HPSE2*-administered *WT* mice (n=3), 63% of 185 ganglion cells were positive for *WPRE3*, and 56% of 210 cells expressed *HPSE2*. In *AAV9/HPSE2*-administered *Mut* mice (n=3 assessed), 74% of 167 ganglion cells were positive for *WPRE3*, and 68% of 191 cells expressed *HPSE2*. Heparanase-2 was sought with an antibody reactive to both human and mouse protein (Figure 4i-l), with consistent findings in three mice examined in each experimental group. The protein was detected in pelvic ganglia of untreated *WT* mice, but only a faint background signal was noted in ganglia of untreated mutant. Heparanase-2 was detected in ganglia of both *WT* and *Mut* mice that had been administered the vector.

**Figure 4.**
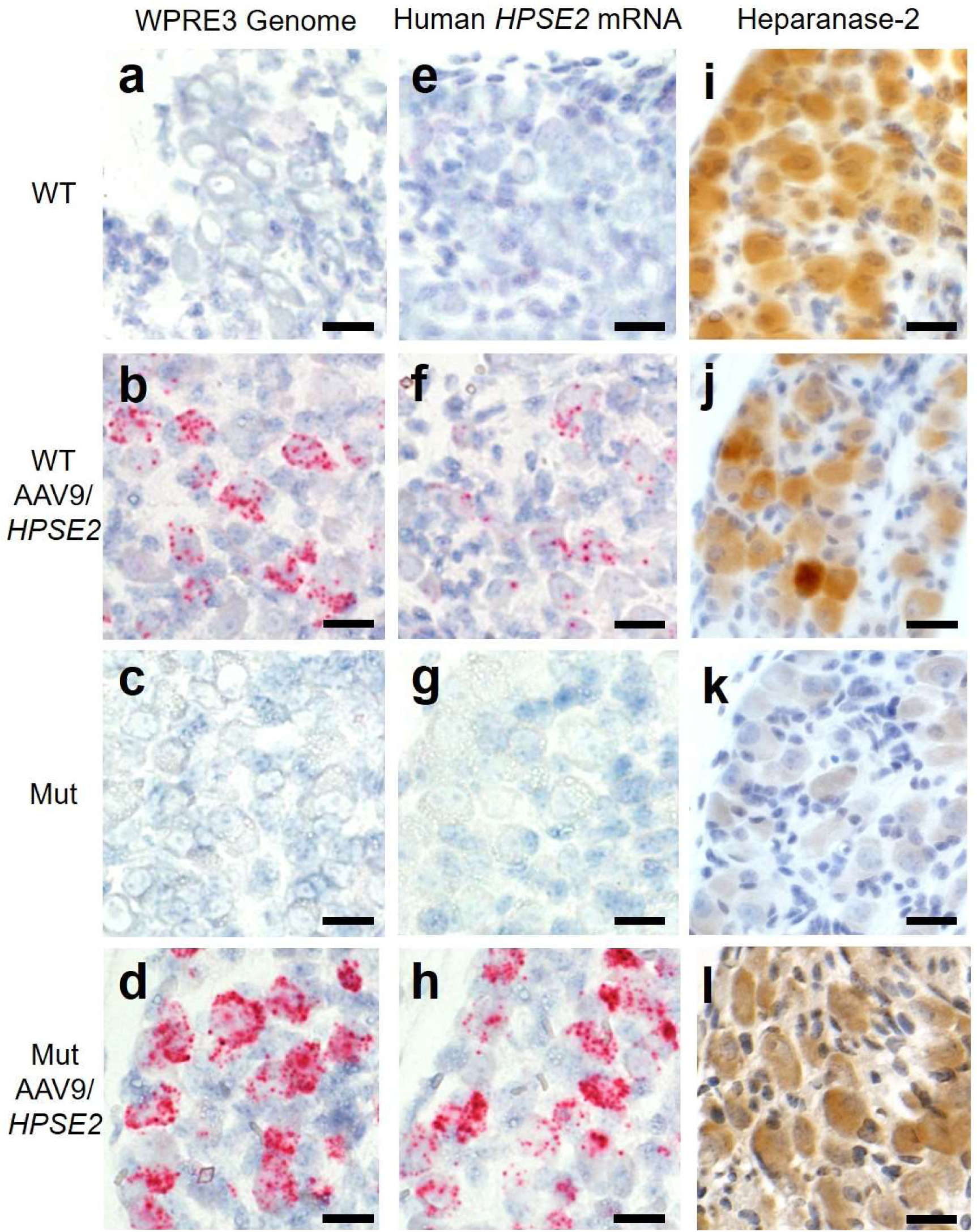
Pelvic ganglia histology in the third week of life. Basescope^TM^ probes were applied for the *WPRE3* genomic sequence (**a-d**) and for human *HPSE2* transcripts (**e-h**). Other sections were reacted with an antibody to heparanase-2 reactive both human and mouse proteins (**i-l**). The four experimental groups were: *WT* mice that were not administered the viral vector (a, e and i); *WT* mice that had been administered 1x10^11^ *AAV9*/*HPSE2* as neonates (**b, f** and **j**); *Mut* mice that were not administered the viral vector (**c, g** and **k**); and *Mut* mice that had been administered 1x10^11^ *AAV9*/*HPSE2* as neonates (**d, h** and **l**). In each group, ganglia from three mice were examined, and representative images are shown. Sections were counterstained with haematoxylin (blue nuclei). Note the absence of Basescope^TM^ signals in both *WT* and *Mut* mice that had not received the viral vector. In contrast, ganglia from *WT* or *Mut* mice that were administered *AAV9/HPSE2* displayed signals for both *WPRE3* and *HPSE2*. Note that the signals appeared prominent in the large cell bodies which are postganglionic neurons; signals were rarely noted in the small support cells between the neural cell bodies. See main text for quantification of signals. Immunostaining for heparanase-2 showed a positive (brown) signal in all groups apart from ganglia from *Mut* mice that had not been administered the viral vector; those cells had only a faint background signal. Bars are 20 μm.

Finally, we determined whether the viral vector transduced cells in the body of the bladder (Figure 5). The vector genome sequence *WPRE3*, and *HPSE2* transcripts, were not detected in the urothelium or lamina propria, the loose tissue directly underneath the urothelium. Within the detrusor muscle layer itself, the large smooth muscle cells were not transduced. However, there were rare small foci of BaseScope^TM^ signal that may represent nerves coursing through the detrusor.

**Figure 5.**
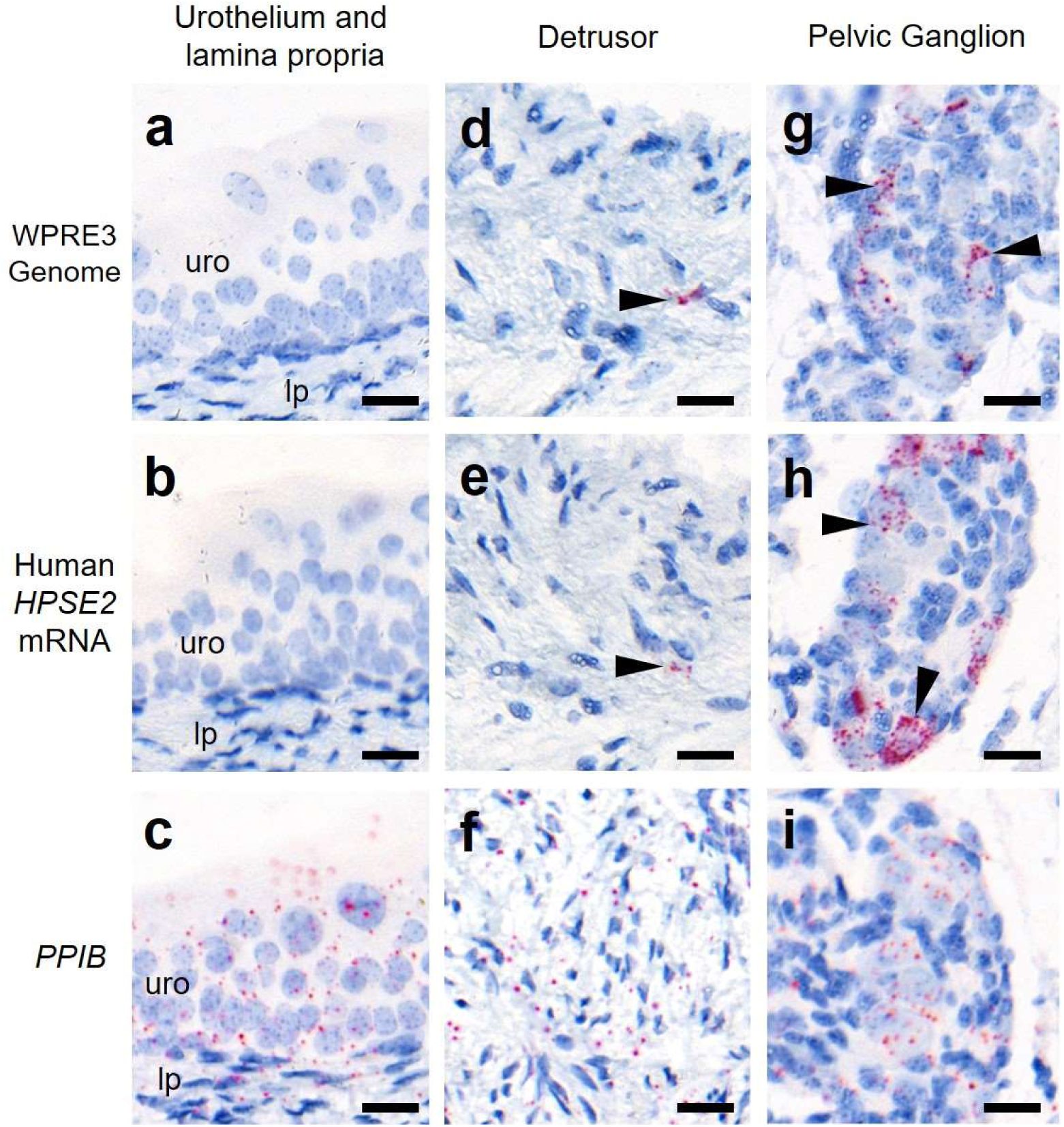
Bladder body histology in the third week of life. Basescope^TM^ ISH analyses of bladder tissue from *AAV9/HPSE2* administered *Hpse2* mutant mice. Images are representative of three mice examined in this manner. Positive signals appear as red dots and nuclei were counterstained with haematoxylin for the bladder urothelium (uro) and lamina propria (lp) (**a-c**), detrusor smooth muscle layer (**d-f**) and pelvic ganglia body (**g-i**). Basescope^TM^ probes were applied for the *WPRE3* genomic sequence (**a**, **d**, **g**), human *HPSE2* transcripts (**b, e, h**) and a positive control transcript, *PPIB* (**c**, **f**, **i**). Note the absence of Basescope^TM^ signals for *WPRE3* and *HPSE2* in the urothelium and lamina propria. There were rare isolated foci of staining in the detrusor layer (indicated by arrow heads in d and e) and abundant staining in the pelvic ganglion (arrowheads in g and h). In contrast, note the widespread expression patterns of the positive control transcript in all tissue types. Scale bars are 20 μm.

The vector genome sequence *WPRE3*, and *HPSE2* transcripts, were also detected in livers of *WT* and *Mut* mice that had been administered the vector (Figure 6a). The average number of vector (red dots per area of liver section) from three livers of *WT* mice injected with *AAV9/HPSE2* were 201/mm^2^ for *WPRE3* and 30/mm^2^ for *HPSE2*. The average values from three *Mut* mice injected with the *AAV9/HPSE2* were 293/mm^2^ for *WPRE3* and was 172/mm^2^ for *HPSE2*. There was no sign of pathological fibrosis in the livers of these mice, as assessed by PSR staining and birefringence quantified under polarised light (Figure 6b and c). Kidneys were also analysed using histology but neither the *WPRE3* sequence nor *HPSE2* transcripts were detected in glomeruli or tubules (Figure 7).

**Figure 6.**
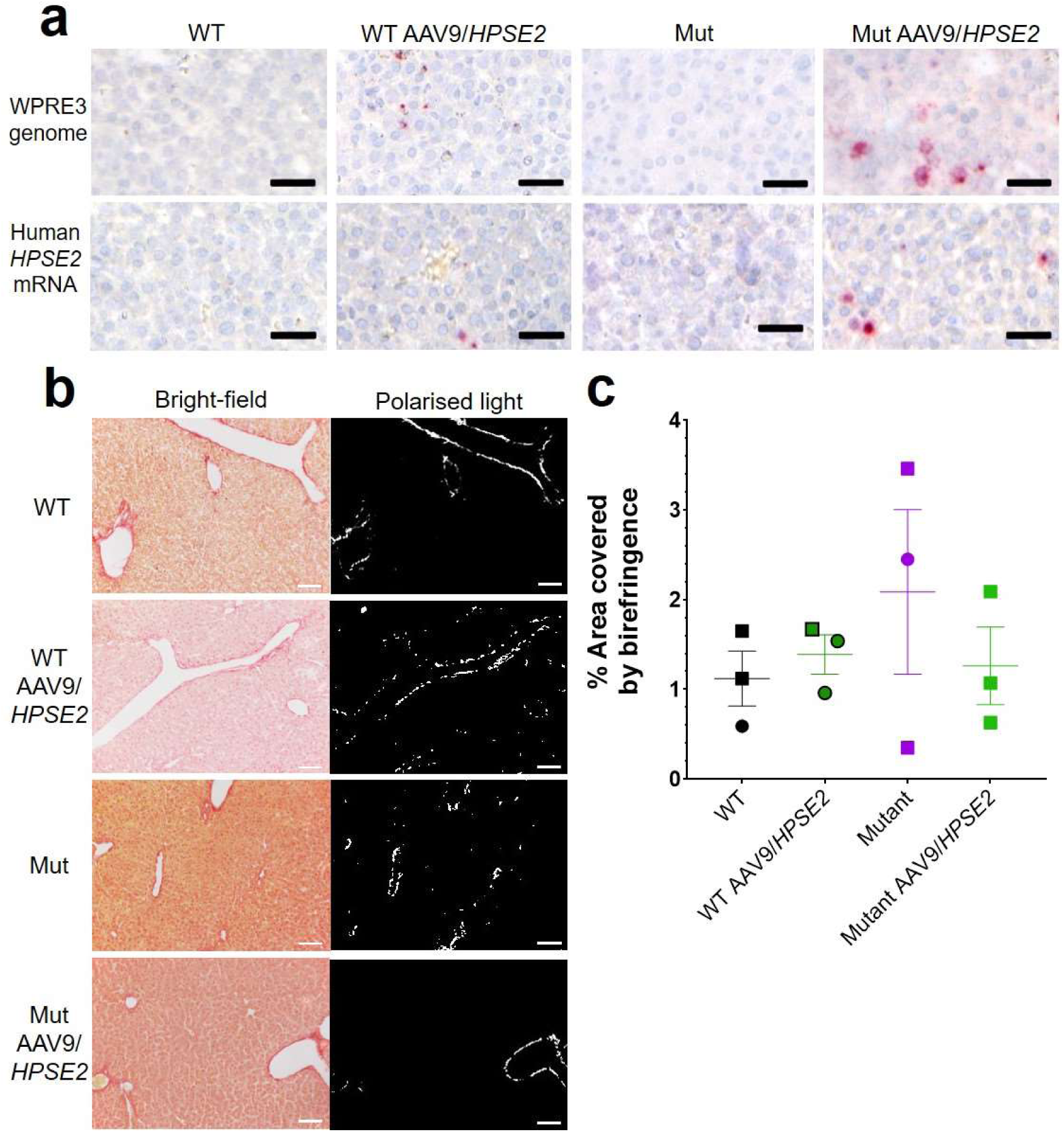
Liver histology in the third week of life. The four experimental groups were: *WT* mice that were not administered the viral vector; *WT* mice that had been administered the viral vector; *Mut* mice that were not administered the viral vector; and *Mut* mice that had been administered *AAV9*/*HPSE2* as neonates. In each group, livers from three mice were examined, and representative images are shown. **a.** Basescope^TM^ ISH analyses of livers, with positive signals appearing as red dots. Nuclei were counterstained with haematoxylin. The vector genome sequence *WPRE3*, and *HPSE2* transcripts, were detected in livers of *WT* and *Mut* mice that had been administered *AAV9/HPSE2*. See main text for quantification. **b.** PSR staining to seek collagen imaged under direct light (left column) and polarised light (right column). There was no significant difference in extent of birefringence between the four groups. Black bars are 30 μm, white are 200 μm.

**Figure 7.**
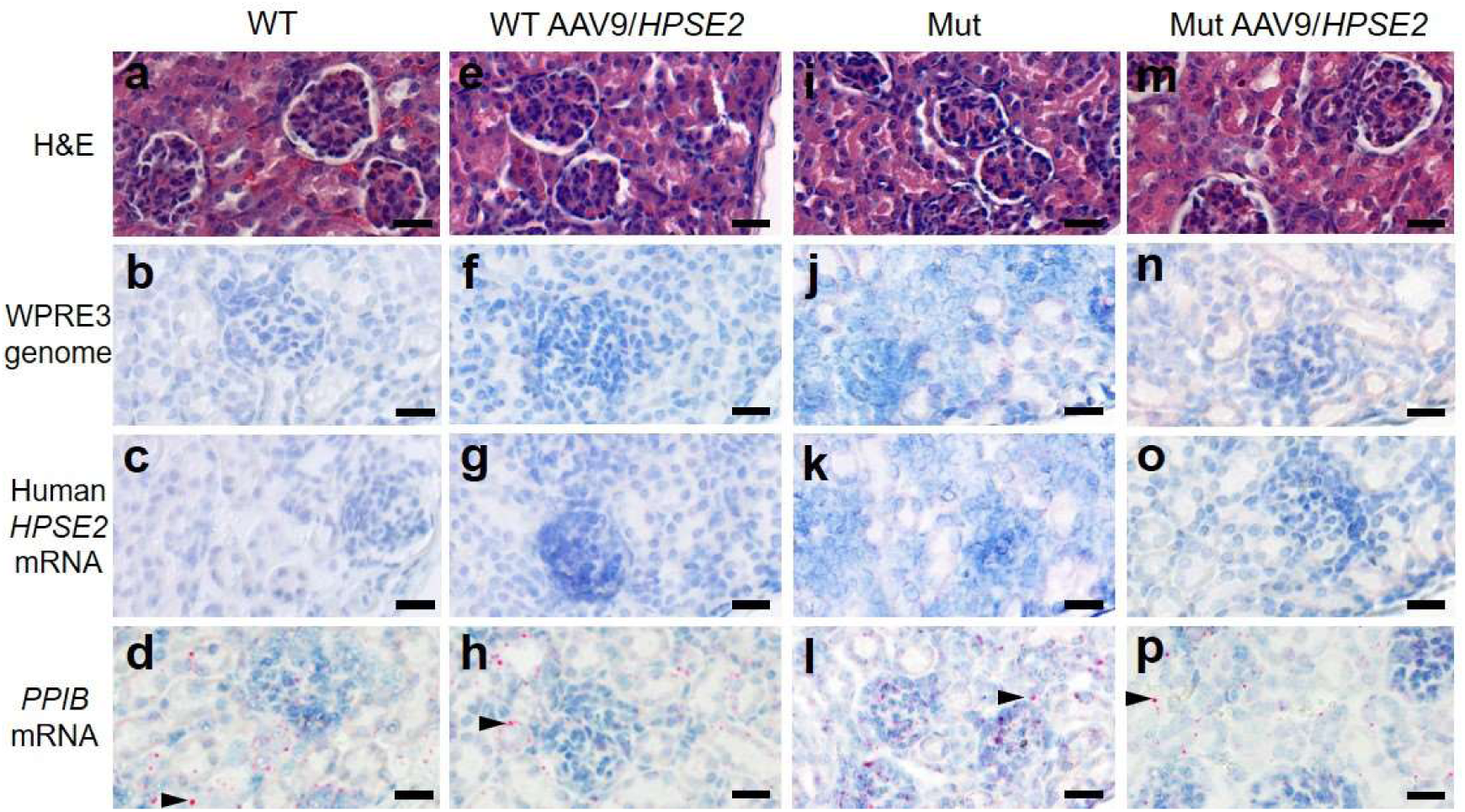
Kidney histology in the third week of life. The four experimental groups were: *WT* mice that were not administered the viral vector (**a-d**); *WT* mice that had been administered 1x10^11^ *AAV9*/*HPSE2* as neonates (**e-h**); *Mut* mice that were not administered the viral vector (**i-l**); and *Mut* mice that had been administered 1x10^11^ *AAV9*/*HPSE2* as neonates (**m-p**). In each group, kidneys from three mice were examined, and representative images are shown. Some sections were stained with haematoxylin and eosin **(a, e, I** and **m)** with similar appearances of glomeruli and tubules in all four groups. Basescope^TM^ probes were applied for the *WPRE3* genomic sequence (**b, f, j** and **n**) and for human *HPSE2* transcripts (**c, g, k** and **o**). Note the absence of Basescope^TM^ signals for *WPRE3* and *HPSE2* in all four groups. In contrast, signals (red dots) were noted after application of a Basescope^TM^ probe for the house-keeping transcript *PPIB* (**d, h, l** and **p**) as shown by arrow heads. Bars are 20 μm.

### *HPSE2* gene transfer ameliorates bladder pathophysiology in *Hpse2* mutant males

We proceeded to undertake *ex vivo* physiology experiments using myography, with isolated bladder outflow tracts and isolated bladder bodies studied separately. No significant differences in contraction amplitudes induced by 50 mM KCl were documented comparing *WT* outflow tracts with *Mut* tissues of mice that had either been administered, or had not been administered, *AAV9/HPSE2* (Figure 8a). Male outflow tracts were pre-contracted with PE, and then subjected to EFS. Nerve-mediated relaxation of pre-contracted outflow tracts had been reported to be impaired in juvenile male *Mut* mice (Manak *et al*, 2020). Accordingly, as expected, in the current study neurogenic relaxation of outflow tracts from untreated *Mut* mice was significantly less than *WT* relaxation (Figure 8b and c). For example, the average *Mut* relaxation at 15 Hz was around a third of the *WT* value. In outflow tracts from *AAV9/HPSE2*-administered *Mut* mice, however, EFS-induced relaxation was no longer significantly different to *WT* controls, and it was three-fold greater than in untreated *Mut* mice (Figure 8b and c).

**Figure 8.**
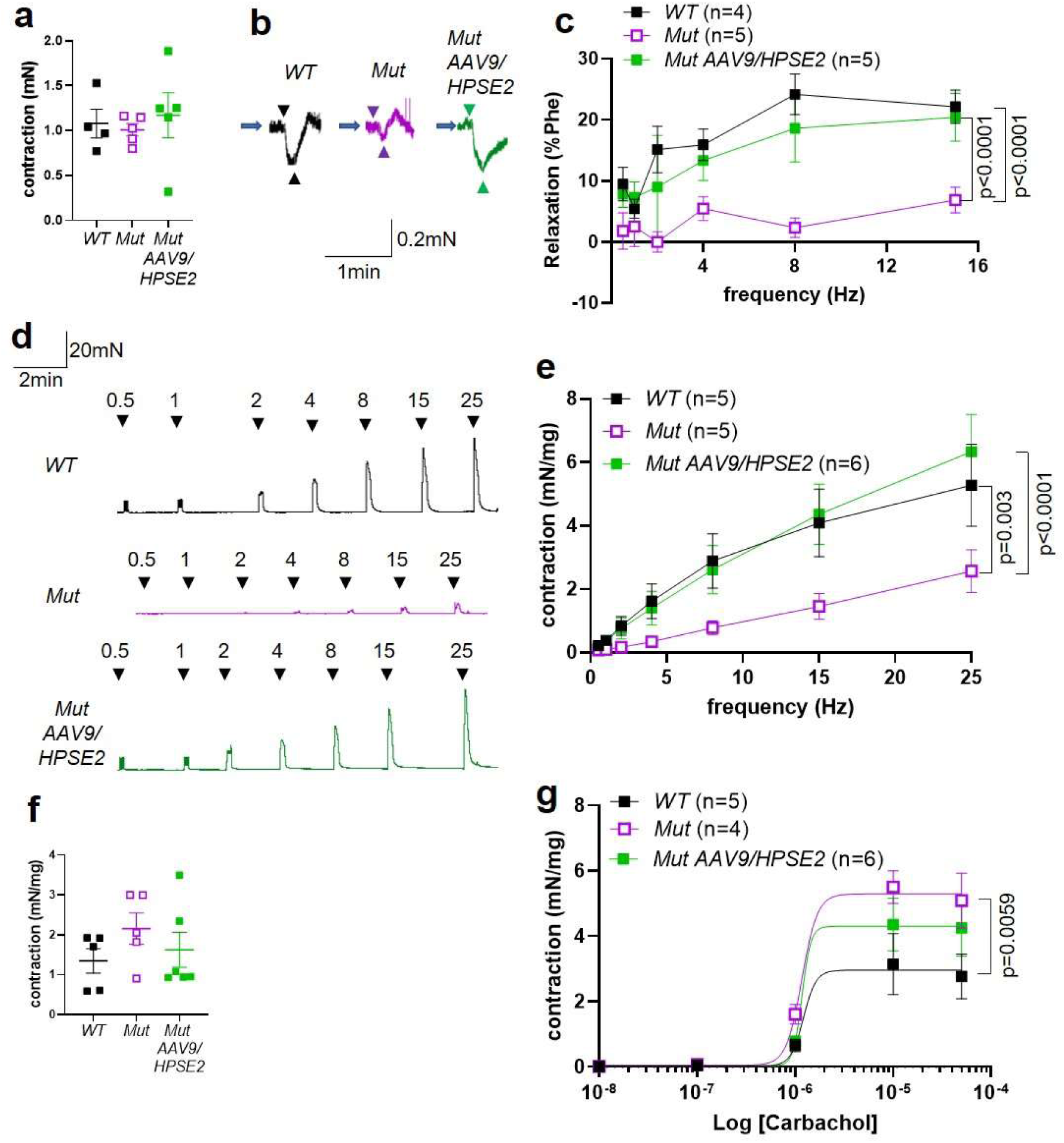
*Ex-vivo* myography in males. **a-c** are bladder outflow tracts, and **d-g** are bladder body rings. **a**. Amplitudes of contraction evoked by 50 mM KCl in male *WT* (n=4), *Mut* (n=5) and *Mut AAV9/HPSE2* (n=5) outflow tracts. **b.** Representative traces of relaxation evoked in *WT*, *Mut* and *Mut AAV9/HPSE2* outflow tracts in response to EFS at 8 Hz, with arrowheads indicating the start and end of stimulation. **c.** Relaxations (mean±SEM) evoked by EFS, plotted as a function of frequency in *WT* (n=4), *Mut* (n=5) and *Mut AAV9/HPSE2* (n=5) outflow tracts. **d.** Representative traces contractions evoked in *WT*, *Mut* and *Mut AAV9/HPSE2* bladder body rings in response to EFS at the frequencies indicated. **e.** Amplitude of contractions (mean±SEM) evoked by EFS in bladders from *WT* (n=5), *Mut* (n=5) and *Mut AAV9/HPSE2* (n=6) mice plotted as function of frequency. **f.** Amplitudes of contraction evoked by 50 mM KCl in *WT* (n=5), *Mut* (n=5) and *Mut AAV9/HPSE2* (n=6) bladders. **g.** Contraction (mean±SEM) of bladder rings from *WT* (n=5), *Mut* (n=4) and *Mut AAV9/HPSE2* (n=6) mice in response to cumulative application of 10 nM-50 µM carbachol, plotted as a function of carbachol concentration. Curves are the best fits of the Hill equation with EC50 = 1.21 µM and Emax= 2.96 mN/mg in *WT* mice compared with EC50 = 1.20 µM and Emax = 5.30 mN/mg for *Mut* mice and EC50 = 1.17 µM and Emax = 4.3 mN/mg in *Mut AAV9/HPSE2* mice. comparing *WT* and *Mut* by 2-way ANOVA with repeated measures.

Next, male bladder body rings were studied using myography (Figure 8d-g). Application of EFS caused detrusor contractions of isolated bladder body rings. In samples from mice that had not received *AAV9/HPSE2* as neonates, contractions were significantly lower in *Mut* than in *WT* preparations (Figure 8d and e). For example, at 25 Hz the average *Mut* value was half of the *WT* value. Strikingly, however, the contractile response to EFS increased around 2.5-fold in *Mut* bladder rings from *AAV9/HPSE2*-administered mice compared with untreated *Mut* mouse bladder rings (Figure 8d and e). Indeed, EFS-induced DSM contractions in treated *Mut* mice were not statistically different to those of *WT* mice (Figure 8e). No significant difference in contraction amplitude to 50 mM KCl was documented comparing *WT* samples with tissues isolated from either vector administered or untreated *Mut* mice (Figure 8f). As previously reported (Manak *et al*, 2020), bladder body contractions in response to cumulative increasing concentrations of carbachol were significantly higher in untreated *Mut* tissues in comparison with *WT* bladder rings (Figure 8g). Bladder body samples from *AAV9/HPSE2*-administered *Mut* mice showed an attenuated hyper-response to carbachol, and values in this treated *Mut* group were no longer significantly different to the *WT* response (Figure 8g).

### *HPSE2* gene transfer ameliorates bladder pathophysiology in *Hpse2* mutant females

UFS affects both males and females (Ochoa, 2004; Newman *et al*, 2023), yet results of *ex vivo* physiology have to date only been reported in tissues from *Hpse2* mutant male mice (Manak *et al*, 2020). Therefore, we proceeded to study outflow tracts and bladder bodies from female *Mut* mice by *ex vivo* myography (Figure 9a-g). No significant difference in contraction amplitude to 50 mM KCl was documented comparing *WT* outflow tracts with *Mut* tissues from mice that had either been administered, or not been administered, *AAV9/HPSE2* (Figure 9a). Female outflow tracts were pre-contracted with vasopressin because they do not respond to PE, the α1 receptor agonist (Grenier *et al*, 2023). In the current study, therefore, vasopressin was used to pre-contract female outflow tracts before applying EFS. Such samples from untreated *Hpse2* female mutant mice showed significantly less relaxation than *WT* outflow tracts (Figure 9b and c), with a striking twenty-fold reduction at 15 Hz. Outflow tract preparations from female *Mut* mice that had been administered *AAV9/HPSE2*, however, displayed EFS-induced relaxation that was not significantly different to *WT* samples (Figure 9 b and c). Indeed the relaxation response of treated *Mut* samples was significantly increased compared with untreated *Mut* mice, with an average twelve-fold increase in relaxation at 15 Hz (Figure 9c).

**Figure 9.**
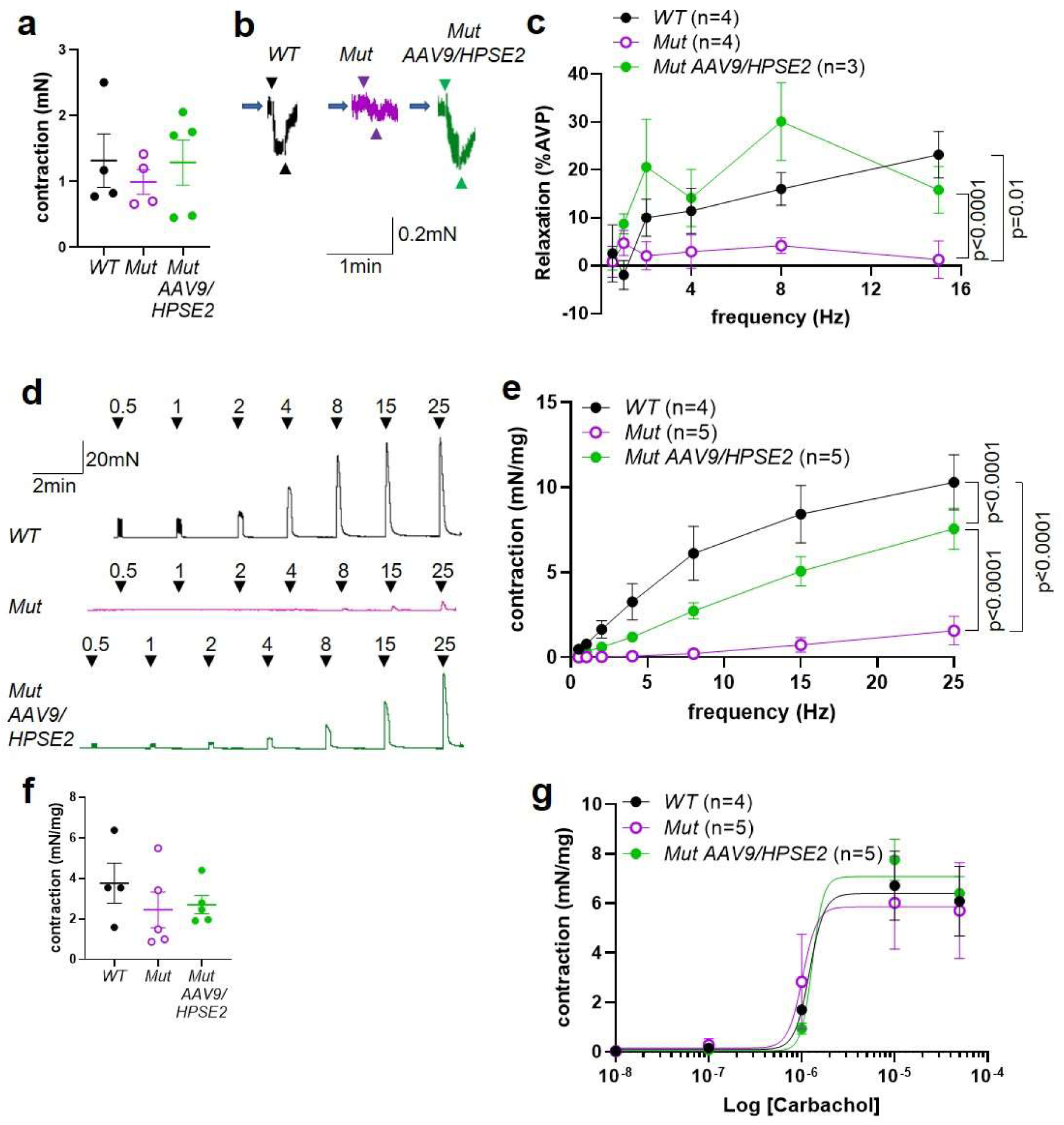
*Ex-vivo* myography in females. **a-c** are outflow tracts, and **d-g** are bladder body rings. **a**. Amplitudes of contraction (mean±SEM) evoked by 50 mM KCl in female *WT* (n=4), *Mut* (n=4) and *Mut AAV9/HPSE2* (n=5) outflow tracts. **b.** Representative traces of relaxation evoked in female *WT*, *Mut* and *Mut AAV9/HPSE2* vasopressin precontracted outflow tracts in response to EFS at 8 Hz. Arrowheads indicate the start and the end of stimulation. **c.** Relaxations (mean±SEM) evoked by EFS, plotted as a function of frequency in female *WT* (n=4), *Mut* (n=4) and *Mut AAV9/HPSE2* (n=3) outflows. **d.** Representative traces of contraction evoked in female *WT*, *Mut* and *Mut AAV9/HPSE2* bladder body rings in response to EFS at the frequencies indicated. **e.** Mean ± SEM Amplitude of contraction (mean±SEM) evoked by EFS in bladder body rings from *WT* (n=4), *Mut* (n=5) and *Mut AAV9/HPSE2* (n=5) mice plotted as function of frequency. **f.** Amplitudes of contraction evoked by 50 mM KCl in female *WT* (n=4), *Mut* (n=5) and *Mut AAV9/HPSE2* (n=5) bladders. **g.** Contraction of bladder rings (mean±SEM) from *WT* (n=4), *Mut* (n=5) and *Mut AAV9/HPSE2* (n=5) mice in response to cumulative application of 10 nM-50 µM carbachol, plotted as a function of carbachol concentration. Curves are the best fits of the Hill equation with EC50 = 1.22 µM and Emax= 6.40 mN/mg in *WT* mice compared with EC50 = 1.02 µM and Emax = 5.86 mN/mg for *Mut* mice and EC50 = 1.30 µM and Emax = 7.08 mN/mg in *Mut AAV9/HPSE2* mice.

Next, female bladder body rings were studied with myography (Figure 9d-g). Applying EFS caused contractions of isolated bladder body rings (Figure 9d and e). In preparations from animals that had not received *AAV9/HPSE2* as neonates, contractions were significantly lower in *Mut* than in *WT* preparations, being reduced seven-fold at 25 Hz. Strikingly, however, the contractile detrusor response to EFS in *Mut* bladder rings from mice that had received *AAV9/HPSE2* was significantly greater than samples from untreated *Mut* mice, with a five-fold increase at 25 Hz. However, the bladder body contractions in *AAV9/HPSE2*-administered *Mut* mice were still significantly smaller than those in *WT* mice (Figure 9e). No significant difference in contraction amplitude to 50 mM KCl was documented comparing the three bladder body groups (Figure 9f). Unlike the enhanced contractile response to carbachol in male *Mut* bladder rings, detailed above, female bladder rings showed similar responses to carbachol in each of the three experimental groups (Figure 9g).

## DISCUSSION

Currently, no treatments exist that target the primary biological disease mechanisms underlying REOLUT disorders. Interventions for UFS are limited and include catheterization to empty the bladder, drugs to modify smooth muscle contractility, and antibiotics for urosepsis (Ochoa, 2004; Newman *et al*, 2023; Osorio *et al*, 2021). Moreover, surgery to refashion the LUT in UFS is either ineffective or even worsens symptoms (Ochoa, 2004). Given that some REOLUT disorders have defined monogenic causes, it has been reasoned that gene therapy might be a future option, but the strategy must first be tested on genetic mouse models that mimic aspects of the human diseases (Lopes *et al*, 2021). The results presented in the current study constitute first steps towards curing UFS, a clinically devastating genetic disease featuring a bladder autonomic neuropathy.

We report several novel observations. First, female *Hpse2* mutant mice have neurogenic defects in outflow tract relaxation and detrusor contraction that are similar to those reported for male *Hpse2* mutant mice (Manak *et al*, 2020). This sex equivalence in the mouse model is an important point because UFS affects both males and females (Newman *et al*, 2023). On the other hand, we observed nuanced differences between the mouse sexes. For example, regarding bladder bodies, females displayed a more profound defect in contractions elicited by EFS, while they lacked the hyper-sensitivity to carbachol shown by males. Second, we have demonstrated that an AAV can be used to transduce pelvic ganglia that are key autonomic neuronal structures in the pathobiology of UFS. Importantly, the neuro-muscular circuitry of the bladder matures over the first three postnatal weeks in murine species (Keast *et al*, 2015). A single dose of 1x10^11^ genome copies of *AAV9/HPSE2* was sufficient to transduce around half of neural cell bodies in these ganglia, when assessed in the third postnatal week. It is possible that the administration of a higher dose of the vector would have resulted in a higher percent of positive cell bodies. Strikingly, heparanase-2 became detectable in pelvic ganglia of treated *Mut* mice. Our third key observation is that the 1x10^11^ dose significantly ameliorated defect in the mutant LUT, despite not every neural cell body being transduced. On autopsy, wild-type mice had empty bladders whereas bladders were uniformly distended in mutant mice, a defect ameliorated by *AAV9/HPSE2* treatment. *AAV9/HPSE2* significantly ameliorated the impaired neurogenic relaxation of outflow tracts and the impaired neurogenic contractility of mutant detrusor smooth muscle found in *Hpse2 Mut* mice. In future, it will be important to study whether the gene transfer strategy might also ameliorate a dyssynergia between bladder and outflow as assessed by *in vivo* cystometry (Ito *et al*, 2018). These complex *in vivo* experiments are, however, beyond the remit of the current work.

Although our data support a neurogenic origin for defects in UFS bladder functionality, we can not yet rule out an additional myogenic component. In a previous study (Stuart *et al*, 2015), however, we quantified smooth muscle actin (*Acta2*) and myosin heavy chain (*Myh11*) transcripts at one day and 14 days after birth and recorded no significant differences between wild-type and *Hpse2* homozygous mutant mice. This suggests that detrusor smooth muscle itself is unchanged in the juvenile period. Moreover, in the current study, we did not detect transduction of *AAV9/HPSE2* into detrusor smooth muscle. A more thorough molecular and structural analysis of the natural history of the detrusor will be an important future work. Similarly, fully elucidating the molecular basis of the UFS neurogenic defect is a key objective. A recent paper has demonstrated the feasibility of single cell RNA sequencing in murine pelvic ganglia cells (Sivori *et al*, 2024) and so, in future, analysis of WT, *Hpse2* mutant and *Hpse2* mutant-treated pelvic ganglia neurons could reveal the fundamental processes that are aberrant in UFS neurons, and are rescued by gene addition treatment.

A feature of the current mouse model, also observed in a different *Hpse2* gene trap mouse (Guo *et al*, 2015), is the poor whole body growth compared with *WT* controls. This becomes more marked during the first month of life and, in the current study, it was not ameliorated by neonatal *AAV9/HPSE2* administration. The cause of the growth impairment has not been established, but, as demonstrated in the current study, is not accompanied by structural kidney damage. Moreover, our unpublished observations found that *Hpse2* mutant pups appear to feed normally, as assessed by finding their stomachs full of mother’s milk. Histological examination of their lungs is normal, excluding aspiration pneumonia. Whatever the cause of the general growth failure, it is notably dissociated from the LUT phenotype, given that *AAV9/HPSE2* administration corrects bladder pathophysiology but not overall growth of the mutant mouse. Of note, a recently published study induced deletion of *Hpse2* in young adult mice (Kayal *et al*, 2023), and this was followed by fatty degeneration of pancreatic acinar cells. Whether the line of *Hpse2* mutant mice used in the current study are born with a similar pancreatic exocrine pathology requires further investigation.

A small subset of individuals with UFS carry biallelic variants of *leucine rich repeats and immunoglobulin like domains 2* (*LRIG2*) (Stuart *et al*, 2013; Grenier *et al*, 2023). LRIG2, like heparanase-2, is immunodetected in pelvic ganglia (Stuart *et al*, 2013; Roberts *et al*, 2019). Moreover, homozygous *Lrig2* mutant mice have abnormal patterns of bladder nerves and display *ex vivo* contractility defects in LUT tissues compatible with neurogenic pathobiology (Roberts *et al*, 2019; Grenier *et al*, 2023). In future, the neonatal gene therapy strategy outlined in the current paper could be applied to ameliorate bladder pathophysiology in *Lrig2* mutant mice. As well as in UFS, functional bladder outflow obstruction occurs in some individuals with prune belly syndrome (Volmar *et al*, 2001) and in megacystis microcolon intestinal hypoperistalsis syndrome (Gosemann *et al*, 2011). Moreover, apart from UFS, functional bladder outflow obstruction can be inherited as a Mendelian trait and some such individuals carry variants in genes other than *HPSE2* or *LRIG2*. These other genes are implicated in the biology of bladder innervation, neuro-muscular transmission, or LUT smooth muscle differentiation (Weber *et al*, 2011; Caubit *et al*, 2016, Woolf *et al*, 2019; Houweling *et al*, 2019; Mann *et al*, 2019; Beaman *et al*, 2019; Hahn *et al*, 2022). Genetic mouse models exist for several of these human diseases and, again, could be used as models for gene therapy.

There are numerous challenges in translating work in mouse models to become effective therapies in humans. First, neonatal mice are immature compared to a newborn person, and the anatomy of their still developing kidneys and LUTs can be compared to those organs in the human fetus in the late second trimester (Lopes & Woolf, 2023). Thus, gene replacement therapy for people with UFS and related genetic REOLUT disorders may need to be given before birth. In fact, UFS can present in the fetal period when ultrasonography reveals a dilated LUT (Newman *et al*, 2023; Grenier *et al*, 2023). The ethics and feasibility of delivering genes and biological therapies to human fetuses are current topics of robust debate (Sagar *et al*, 2020; Mimoun *et al*, 2023) and, with advances in early detection and therapies themselves, may become established in coming years. Second, there are concerns about possible side-effects of administering AAV vectors (Srivastava *et al*, 2023). Factoring weight for weight, the dose of *AAV9* vector administered to the *Hpse2* mutant mice in the current study is of a similar order of magnitude as those used to treat babies with spinal muscular atrophy (Mendell *et al*, 2017). In that report there was evidence of liver damage, as assessed by raised blood transaminases in a subset of treated patients. In the current mouse study, we found that the viral vector transduced livers, while kidneys were resistant to transduction. As assessed by histological analyses to seek fibrosis, livers from *AAV9/HPSE2* treated mice appeared normal. We cannot, however, exclude transient liver damage, which could be assessed by alterations in blood transaminases. The fact that transduced livers expressed *HPSE2* transcripts, and the observation that heparanse-2 is a secreted protein (Levy-Adam *et al*, 2010; Beaman *et al*, 2022), raises the possibility that livers of treated mice may act as a factory to produce heparanase-2 that then circulated and had beneficial effects on the LUT, akin to gene therapy to replace coagulation factors missing in haemophilia (Chowdary *et al*, 2022).

In summary, we have used a viral vector-mediated gene supplementation approach to ameliorate tissue-level neurophysiological defects in UFS mouse bladders. This advance provides a proof-of-principle to act as a paradigm for treating other genetic mouse models of REOLUT disorders. In the longer term, application to humans may need to consider fetal therapy, for reasons detailed above.

## METHODS

### Experimental strategy

In a first set of experiments to explore the feasibility of transducing pelvic ganglia, neonatal homozygous wild-type (*WT*), mice were intravenously administered the *AAV9*/*HPSE2* vector (described below). These mice were observed until they were young adults when their internal organs were collected to seek evidence of transgene transduction and expression. Next, we determined whether neonatal administration of *AAV9*/*HPSE2* to homozygous *Hpse2* mutant mice, hereafter called *Mut*, would ameliorate *ex vivo* bladder physiological defects that had been described in juvenile *Hpse2* mutants (Manak *et al*, 2020). The third postnatal week was used as the end point, because *Hpse2 Mut* mice subsequently fail to thrive (Stuart *et al*, 2015) and *ex vivo* physiological bladder aberrations have been characterised in *Mut* mice of a similar age (Manak *et al*, 2020).

### The viral vector

The custom made (Vector Biolabs) *AAV9* vector carried the full-length coding sequence for human *HPSE2* and the broadly active synthetic *CMV early enhancer/chicken Actb* (*CAG*) promoter. This promoter has been used successfully in human gene therapy for spinal muscular atrophy (Mendell *et al*., 2017). The construct also contained the woodchuck hepatitis virus post-transcriptional regulatory element (*WPRE3*), known to create a tertiary structure that increases mRNA stability and thus potentially enhancing biological effects of genes delivered by viral vectors (Loeb *et al*, 1999; Lee *et al*, 2005), and flanking *AAV*2 inverted terminal repeat (*ITR*) sequences.

### Experimental mice

Mouse studies were performed under UK Home Office project licenses PAFFC144F and PP1688221. Experiments were undertaken mindful of ARRIVE animal research guidelines (Percie du Sert *et al*, 2020). C57BL/6 strain mice were maintained in a 12-hour light/dark cycle in the Biological Services Facility of The University of Manchester. The mutant allele has the retroviral gene trap VICTR4829 vector inserted into intron 6 of *Hpse2* (Stuart *et al*, 2015) (mouse accession NM_001081257, and Omnibank clone OST441123). The mutation is predicted to splice exon 6 to a *Neo* cassette and generate a premature stop codon. Accordingly, homozygous mutant mouse bladder tissue contains a truncated *Hpse2* transcript and lacks full length *Hpse2* transcripts extending beyond the trap (Stuart *et al*, 2015). Mating heterozygous parents leads to the birth of *WT*, heterozygous, and *Mut* offspring in the expected Mendelian ratio (Stuart *et al*, 2015). In the first month after birth *Mut* mice gain less weight than their *WT* littermates (Stuart *et al*, 2015) and, because they later fail to thrive, *Mut* mice are culled around a month of age before they become overtly unwell. Application of tattoo ink paste (Ketchum, Canada) on paws of neonates (i.e., in the first day after birth) was used to identify individual mice, and same-day genotyping was undertaken. This was carried out using a common forward primer (5′CCAGCCCTAATGCAATTACC3′) and two reverse primers, one (5′TGAGCACTCACTTAAAAGGAC3′) for the *WT* allele and the other (5′ATAAACCCTCTTGCAGTTGCA3′) for the gene trap allele. Neonates underwent general anaesthesia with 4% isoflurane, and *AAV9/HPSE2* suspended in up to 20 μl of sterile phosphate buffered saline (PBS) was injected into the temporal vein. This procedure took around two minutes, after which the baby mice were returned to their mother. Control neonates received either vehicle-only (PBS) injection or were not injected. At the end of the study, mice underwent Schedule 1 killing by inhalation exposure to a rising concentration of carbon dioxide followed by exsanguination.

### Bladder outflow tract and bladder body myography

*Ex vivo* myography and its interpretation were undertaken using methodology detailed in previous studies (Manak *et al*, 2020; Grenier *et al*, 2023). Functional smooth muscle defects of both the outflow tract and bladder body were previously characterised in juvenile *Hpse2 Mut* mice (Manak *et al*, 2020), so mice of the same age were used here. Outflow tracts, which contained SM but not the external sphincter, were separated from the bladder body by dissection. Intact outflow tracts or full thickness rings from the mid-portion of bladder bodies, containing detrusor smooth muscle, were mounted on pins in myography chambers (Danish Myo Technology, Hinnerup, Denmark) containing physiological salt solution (PSS) at 37°C. The PSS contained 122 mM NaCl, 5 mM KCl, 10 mM N-2-hydroxyethylpiperazine-N’-2-ethanesulfonic acid (HEPES), 0.5 mM KH_2_PO_4_, 1 mM MgCl_2_, 5 mM D-glucose and 1.8mM CaCl_2_ adjusted to pH 7.3 with NaOH.

The contractility of outflow tract and bladder body preparations was tested by applying 50 mM KCl to directly depolarise SM and stimulate Ca^2+^ influx through voltage gated Ca^2+^ channels. Sympathetic stimulation of α-1 adrenergic receptors mediates urethral contraction in male mice and primates but has little effect in females (Alexandre *et al*, 2017). The α-1 adrenergic receptor agonist, phenylephrine (PE, 1 μM), was therefore used to contract male outflow tracts before applying electrical field stimulation (EFS; 1 ms pulses of 80 V at 0.5–15 Hz for 10 s) to induce nerve-mediated relaxation. Female outflow tracts were instead pre-contracted with vasopressin (10 nM) as previously detailed (Grenier *et al*, 2023), because they are known to express contractile arginine-vasopressin (AVP) receptors (Zeng *et al*, 2015). Bladder body rings were subjected to EFS (1 ms pulses of 80V at 0.5–25 Hz for 10 s) to induce frequency-dependent detrusor contractions. As these contractions are primarily mediated by acetylcholine, through its release from parasympathetic nerves and binding to muscarinic receptors (Matsui *et al*, 2002), we also measured the degree of bladder body contractions evoked directly by the muscarinic agonist, carbachol (10 nM – 50 μM).

### Histology

Samples of bladder body, liver, kidney and pelvic ganglia flanking the base of the mouse bladder (Keast *et al*, 2015; Roberts *et al*, 2019) were removed immediately after death and prepared for histology. Tissues were fixed in 4% paraformaldehyde, paraffin-embedded and sectioned at 5 μm for bladders and kidneys, or 6 μm for livers. After dewaxing and rehydrating tissue sections, BaseScope^TM^ probes (ACDBio, Newark, CA, USA) were applied. *In situ* hybridisation (ISH) was undertaken, using generic methodology as described (Lopes *et al*, 2019). To seek expression of transduced *HPSE2*, BA-Hs-*HPSE2*-3zz-st designed against a sequence in bases 973-1102 in exons 5–7 was used. Other sections were probed for the widely expressed *PPIB* transcript encoding peptidylprolyl isomerase B (BaseScope™ Positive Control ProbeMouse (Mm)-PPIB-3ZZ 701071). As a negative control, a probe for bacterial *dapB* (BaseScope™ Negative Control ProbeDapB-3ZZ 701011) was used (not shown). We also used a BaseScope^TM^ ISH probe to seek genomic *WPRE3* delivered by the viral vector (cat. 882281). BaseScope^TM^ detection reagent Kit v2-RED was used following the manufacturer’s instructions, with positive signals appearing as red dots within cells. Gill’s haematoxylin was used as a counterstain. Images were acquired on a 3D-Histech Pannoramic-250 microscope slide-scanner using a *20x/ 0.80 Plan Apochromat* (Zeiss). Snapshots of the slide-scans were taken using the Case Viewer software (3D-Histech). Higher powered images were acquired on an Olympus BX63 upright microscope using a DP80 camera (Olympus) through CellSens Dimension v1.16 software (Olympus). For immunohistochemistry, after dewaxing and rehydration, endogenous peroxidase was quenched with hydrogen peroxide. Slides were microwaved and then cooled at room temperature for 20 min in antigen-retrieval solution (10 mM sodium citrate, pH 6.0). A primary antibody raised in rabbit against heparanase-2 (Abcepta AP12994c; 1:400) was applied. The immunogen was a KLH-conjugated synthetic peptide between 451-480 amino acids in the C-terminal region of human heparanase-2, predicted to cross-react with mouse heparanase-2. The primary antibody was prepared in PBS Triton-X-100 (0.1%) and 3% serum and incubated overnight at 4 °C. It was reacted with a second antibody and the positive brown signal generated with DAB peroxidase-based methodology (Vector Laboratories, SK4100). Sections were counterstained with haematoxylin alone, or haematoxylin and eosin. Omission of the primary antibody was used as a negative control. Some liver sections were also reacted with 1% picrosirius red (PSR) solution (diluted in 1.3% picric acid) for one hour. Birefringent collagen was visualised under cross-polarised light and the percentage of area covered by birefringence measured as described (Hindi *et al*, 2021).

### Statistics

Data organised in columns are expressed as mean±SEM and plotted and analysed using the GraphPad Prism 8 software. The Shapiro-Wilk test was used to assess normality of data distributions. If the values in two sets of data passed this test and they had the same variance, a Student t-test was used to compare them. If the variance was different, a t-test with Welch’s correction was used. If data did not pass the normality test, a non-parametric Mann-Whitney test was used. Groups of data with repeated measurements (e.g. at different stimulation frequencies or concentration) were compared using 2-way ANOVA with repeated measures. An F-test was used to assess the likelihood that independent data sets forming concentration-response relationships were adequately fit by a single curve.

## DISCLOSURE

The authors declare no conflicts of interest.

## Acknowledgements

We acknowledge receiving research funding from: Medical Research Council (project grant MR/L002744/1 to ASW and WGN; project grant MR/T016809/1 to ASW, NAR, and FML; and Doctoral Training Programme studentship to BWJ); Kidney Research United Kingdom (project grant Paed_RP/002/20190925 to WGN, GMB, and ASW; and Paed_RP/005/20190925 to NAR and ASW); Newlife Foundation (project grants 15-15/03 and 15-16/06 to WGN and ASW); the Manchester NIHR Biomedical Research Centre (IS-BRC-1215-20007 to WGN); and Kidneys for Life (start-up grant 2018 to NAR, and project grant to NAR 2023); a LifeArc Pathfinder Award (to NAR, ASW, CG, WGN); and an MRC-NIHR UK Rare Disease Research Platform MR/Y008340/1 to NAR, ASW, WGN. We thank the Manchester NIHR Biomedical Research Centre Rare Diseases theme, and the Manchester Rare Condition Centre for support.

